# Ligation-assisted target recycling for DNA nanoswitch biosensors

**DOI:** 10.64898/2026.05.15.725157

**Authors:** Vinod Morya, Andrew Hayden, Mona Zeghal, Jibin Abraham Punnoose, Ken Halvorsen

**Affiliations:** The RNA Institute, University at Albany, State University of New York, Albany, NY, 12222 USA; Department of Chemistry, State University of New York, New Paltz, NY, 12561 USA

## Abstract

Conformationally responsive DNA nanoswitches have previously been developed and validated for a variety of biosensing applications including detection of DNA, microRNA, and viral RNA/DNA. Here we develop new methodology for enhancing the sensitivity of DNA-based sensing by recycling a fixed number of targets for repeated reuse. We achieved target-dependent enzymatic ligation of looped nanoswitches and showed that subsequent removal of target does not affect the ligated loop. Through cyclic annealing, ligation, and target removal, we can linearly control signal amplification up to hundreds of cycles. This method adds an important new capability for low abundance targets without the need for target amplification.

## Introduction

Sensitive and specific detection of nucleic acids is central to many applications in modern biology, medicine, and biotechnology. From early disease diagnostics to environmental monitoring and fundamental studies of gene regulation, biosensing technologies have evolved to meet the increasing demand for rapid, accurate, and low-cost detection of molecular targets. Among these targets, short nucleic acid sequences such as microRNAs (miRNAs), viral RNA fragments, and cell-free DNA have emerged as critical biomarkers, often present at low abundance and within complex biological environments^1–4^. Detecting such targets remains challenging due to their small size, sequence similarity to off-target molecules, and typically low copy numbers. These constraints have motivated the development of innovative biosensing strategies that combine molecular specificity with signal amplification mechanisms^1,5–7^. DNA nanotechnology offers a powerful framework for engineering such biosensors. Leveraging the predictable base-pairing of nucleic acids, DNA can be programmed to self-assemble into well-defined nanostructures with precise geometries and functionalities^8,9^. Over the past two decades, advances in this field have enabled the construction of dynamic DNA devices capable of responding to molecular inputs through conformational changes, strand displacement reactions, and enzymatic processes. These capabilities have been harnessed to create a diverse array of biosensors that translate molecular recognition events into measurable outputs, including fluorescence, colorimetric signals, and mechanical transitions^10–12^.

One type of DNA-based biosensors are conformationally responsive DNA nanoswitches. DNA nanoswitches are long, duplex DNA constructs assembled from ssDNA scaffolds that are engineered with programmable detector regions^13^. Upon binding to a target molecule, these detector regions bring distant parts of the DNA strand into proximity, inducing a topological transition from a linear to a looped state. This conformational change can be readily detected using standard agarose gel electrophoresis, where the looped and linear forms migrate separately. The simplicity of this readout, combined with the inherent specificity of nucleic acid hybridization, makes DNA nanoswitches an attractive platform for low-cost and accessible diagnostics. Since their initial development, DNA nanoswitches have been applied to detect a variety of biomolecular targets, including nucleic acids^14^, proteins^15^, small molecules^13^ and RNA modifications^16^. In the context of nucleic acid detection, nanoswitches offer several advantages: they are enzyme-free in their basic configuration, require minimal instrumentation, and can achieve specificity up to single base-pair mismatch through careful design of detector strands^17^. Furthermore, the modularity of the platform allows for multiplexed detection by encoding different loop sizes corresponding to distinct targets^18^. These features position DNA nanoswitches as a promising alternative to conventional amplification-based techniques such as polymerase chain reaction (PCR), particularly in resource-limited settings.

One of the key constraints in DNA nanoswitch sensing is a 1:1 stoichiometry between target and signal. Each DNA nanoswitch is looped by one target molecule and only remains looped in the presence of the target. In the absence of an amplification mechanism, the detection limit is set by the number of target molecules. Strategies such as increasing detector affinity, optimizing buffer conditions, and extending incubation times can often improve binding and detection, but they do not fundamentally overcome the 1:1 stoichiometry limitation. For long nucleic acid targets such as viral RNA, one strategy has been to fragment the target into distinct smaller targets, overcoming the 1:1 stoichiometry limitation by physically producing more targets^19^. However, this strategy is limited to long nucleic acids and not readily generalizable to shorter ones.

To address the inherent limitations of DNA nanoswitch-based detection with a fixed population of target molecules, we consider a ligation-based approach for signal amplification that could break the 1:1 stoichiometry limitation. In particular, ligation could enable permanent preservation of a looped nanoswitch even after the target is released to trigger the conversion of another nanoswitch in a bind-ligate-unbind cycle. Splint ligation, in which a target nucleic acid serves as a template to align two adjacent probe strands for their enzymatic joining, provides both high specificity and a direct route to signal generation. The conceptual compatibility between DNA nanoswitches and splint ligation presents an intriguing opportunity to integrate these two approaches. In this combined strategy that we call “ligation-assisted target recycling”, the target molecule would 1) hybridize with the detector strands to trigger the nanoswitch to loop, 2) facilitate splint ligation to covalently secure the loop, and 3) release after ligation to enable multiple ligation events in a recurring cycle. (**Figure 1a**)

**Figure 1.**
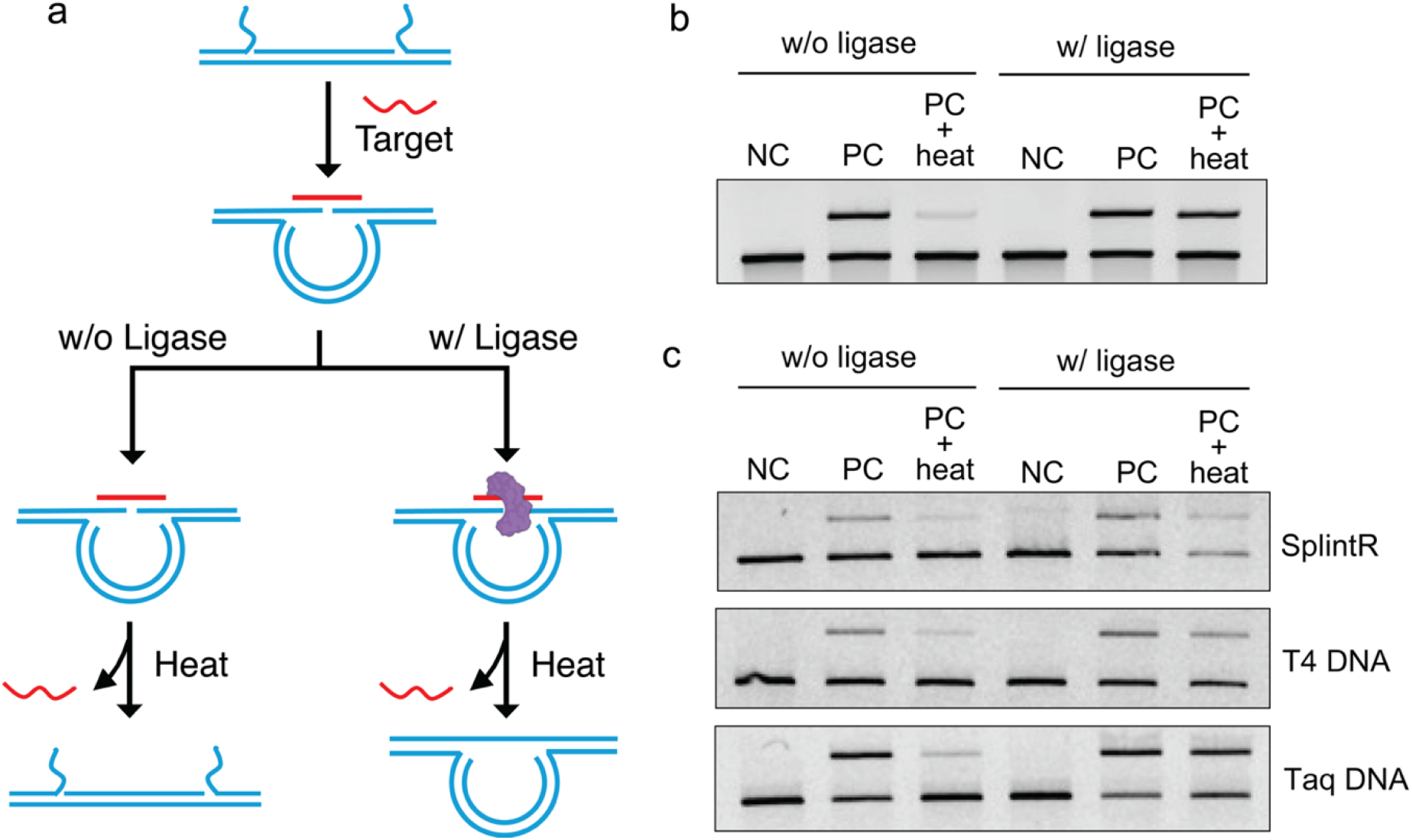
a) Schematic of ligation mediated irreversible looping and b) gel electrophoresis showing irreversible nature of the ligated loop. C) Testing different DNA ligases for ligating the looped nanoswitches.

To develop and validate this strategy to enable target recycling, we first established a ligation scheme that proceeds selectively in the presence of the target and results in loop formation. As a model system, we used our previously developed DNA nanoswitch for the 22 nt DNA sequence (5’-TGAGGTAGTAGGTTGTGTGGTT-3’) corresponding to microRNA let-7b^17^. We matched two 11 nt ssDNA detector regions each complementary to one half of the target. When the nanoswitches are reacted with the target, the loop is forced to close and the looped nanoswitch can be visualized using gel electrophoresis.

Many ligation reactions can repair nicked DNA in a duplex when there is a 3’ OH and a 5’ Phosphate at the nick site. The looped junction of the nanoswitch resembles a double stranded duplex with a central “nick” where the 3’ end of one detector is brought near the 5’ end of another detector. To enable ligation of the nanoswitch at this junction, we integrated a 5’ phosphate into the relevant detector strand. To test the ligation, the nanoswitch was first incubated with the target to induce loop formation, followed by a heating step to dissociate the target (**Figure 1b**). In the absence of ligation, removal of the target resulted in a reversal to the linear state, confirming removal of the target. In contrast, when ligation occurred, the looped state persisted after target removal, consistent with covalent sealing of the junction.

Having successfully established nanoswitch ligation, we next focused on identifying conditions to enable efficient target recycling. Temperature provides an ideal way to enable rapid and controllable target release with simple cycling but requires thermostable ligases that remain active across a broader temperature range. We evaluated the binding behavior of the nanoswitch across a range of temperatures to identify suitable annealing and removal conditions. We observed that target binding occurs efficiently below ∼40 °C, with maximal looping at 35 °C (**Figure S1a**). We then examined the temperature and duration required for effective target dissociation (**Figure S1b**,**c**) and found that heating to elevated temperatures for just ∼20 seconds enabled efficient target removal. Considering that the melting temperatures of the detector-target duplexes are approximately 55 °C, we selected 35 °C for annealing and 65 °C for denaturation in subsequent experiments. To prevent dissociation of the detector oligos from the nanoswitch during the target removal step, we evaluated different backbone oligo lengths of 30 nt, 40 nt, and 60 nt (**Figure S2**). We found that the detector oligo containing a 60 nt backbone provided the greatest stability and maintained loop integrity even at 70 °C.

Next, we evaluated several commercially available DNA ligases for their ability to catalyze ligation at 30°C, a temperature compatible with the operating range of most ligases. Specifically, we tested SplintR, Taq, and T4 ligases for activity under these conditions (**Figure 1c**). Although all three enzymes exhibit some ligation activity near this temperature, Taq ligase demonstrated the most robust performance and retained activity across a broader temperature range, making it well-suited for thermal cycling in our system.

Having a validated ligase and optimized temperatures for binding, ligation, and release, we sought to demonstrate signal enhancement enabled by target recycling. A cycling reaction illustrates the process and the parameters chosen for this work (**figure 2a,b**). We performed temperature cycling between 35 °C and 65 °C in the presence of ligase and a limiting concentration of target DNA, varying the number of cycles and comparing the results to reactions incubated at a constant temperature for the same total duration. We observed an initially linearly increasing trend that plateaus at higher cycles when unlooped substrate becomes rarer (**Figure 2c**). The final signal exceeded that obtained from any constant-temperature incubation over the same time period (**Figure S3)**.

**Figure 2.**
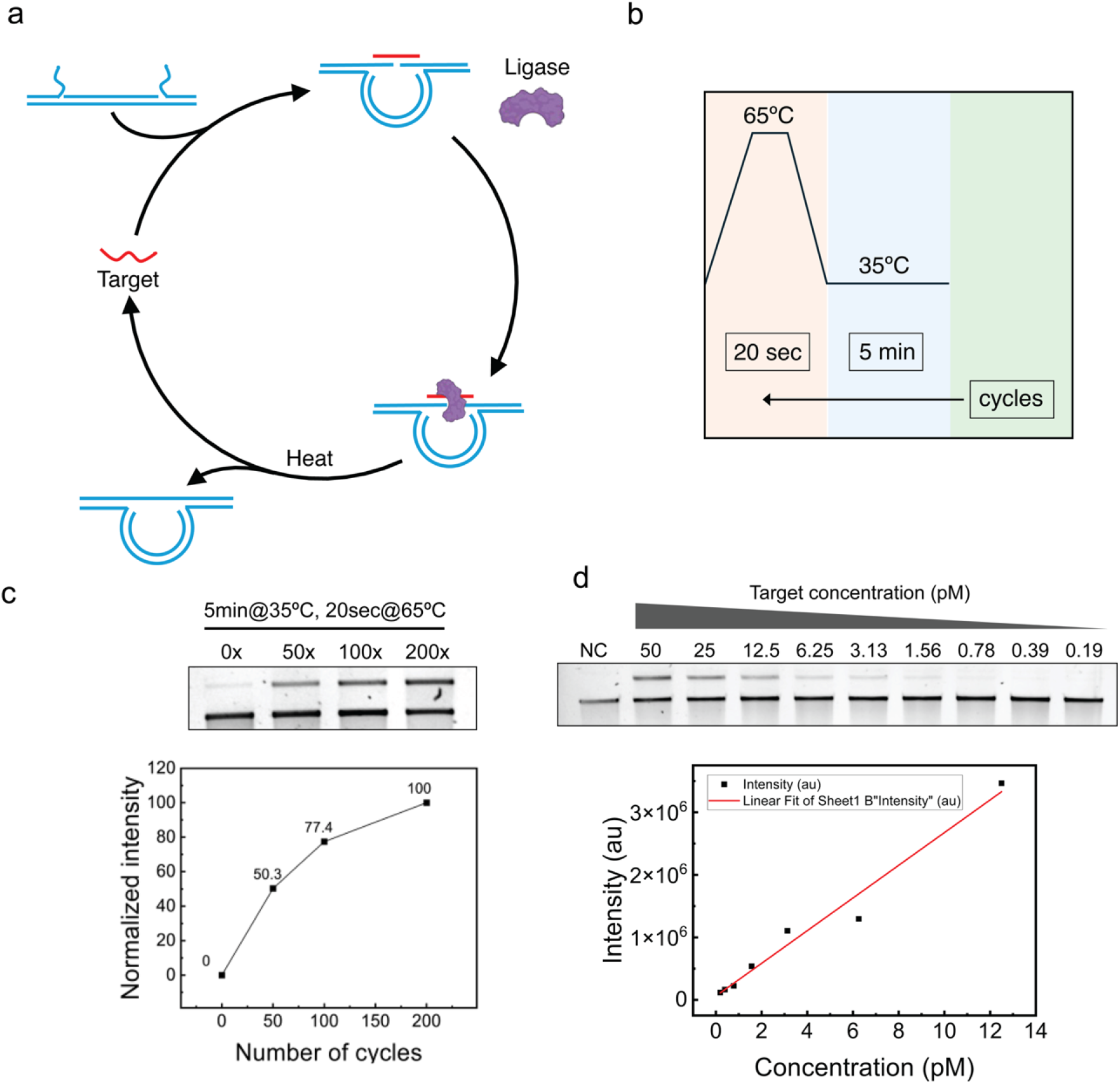
Schematic of a) target recycling reaction: The target initiates loop formation at the annealing temperature, and the ligase subsequently seals the nick between the detector oligos, resulting in irreversible loop formation. A subsequent heating step removes the target, allowing it to participate in the next reaction cycle. b) temperature cycles for target removal and annealing. c) Testing the cycle number depended looping efficiency with a constant 25 pM target concentration. d) Sensitivity test for let-7b DNA and limit of detection assay based on the intensity values.

With the optimized conditions for annealing temperature and time, as well as target removal temperature and duration, we next evaluated the effect of ligase concentration on the reaction. We tested 0 U, 0.4 U, 4 U, and 10 U of Taq ligase in a 10 µL reaction volume and observed that loop formation reached saturation at 4 U of ligase (**Figure S4**). To assess nanoswitch sensitivity with the ligation assisted target recycling, we tested serial dilutions of the target with a fixed 50 cycles (Figure 2d & S5). We observed visible detection down to 190 fM, and calculated the limit of detection as 27 fM. These results demonstrate for the first time that DNA nanoswitches can be paired with a ligation step to enable covalent closure of the loop. This preserves the signal even when the target has become unbound, allowing a mechanism for target recycling to amplify signal with a fixed amount of target. We clearly show signal increase with cycle number, and exceed our normal detection limits for DNA nanoswitches.

The work was aimed at providing proof of concept for this approach, and it is clear that some limitations exist, and further optimization will almost certainly be required. The strategy requires temperature cycling, with annealing and release temperatures that may be different for different targets. We had also considered an alternative strategy of toehold-mediated strand displacement, which could offer isothermal operation, but presents practical challenges including need for additional strands and relatively slow kinetics. Inefficiencies may also be encountered in the amplification strategy, since ligated targets become an increasingly large portion of nanoswitches and may even preferentially bind targets due to the increased binding energy.

Future work will focus on establishing the approach for RNA targets, which seems like a straightforward extension but carries some serious challenges. Most notably, RNA targets require a different ligase to repair DNA nicks in a DNA/RNA hybrid while still maintaining thermostability and ligating around 30-40 ºC. Further improvements in the method would be refinement of reaction times for faster cycling, as well as adaptations for binding of weaker targets including shorter nucleic acid fragments. Aspirationally, this method could pave the way to direct single-molecule detection of nucleic acids, especially if thousands of cycles could be implemented. While the general approach described here is focused on our DNA nanoswitches, it may also find utility in other DNA-based sensors with a similar detection mechanism.

## Materials and Methods

### Reagents

DNA oligos were procured from Integrated DNA Technologies (IDT), US, the unmodified oligos were with standard desalting and the 5’ phosphate modified oligos are PAGE purified. Detailed list of oligos is provided in the Supplementary Information Table 1. The Taq DNA ligase (#M0208), SplintR ligase (#M0375) and T4 DNA ligase (#M0202) with their respective reaction buffers were purchased from New England Biolabs (NEB), US. All the agarose gels presented in manuscript were prepared from Sigma-Aldrich agarose, Bioreagent low EEO (Cat. No. A9539) in freshly prepared 0.2 µm filter sterilized 0.5X tris borate EDTA buffer (TBE).

### Nanoswitch assembly and purification

The nanoswitch assembly was carried out as previously reported protocol^20,21^, that starts with linearization of circular M13 ssDNA (NEB # N4040). The linearized M13 ssDNA was mixed with 10-fold excess of backbone oligos, set of variable strands and a pair of detector stands with filler stands if required. The annealing process was done in a Bio-Rad T100 Thermal Cycler by heating the samples to 90ºC and gradually colling (1ºC per minute) till 20ºC. After annealing, the samples were purified with liquid chromatography.

### Nanoswitch reactions without ligase

The nanoswitch reactions were carried out in a total volume of 10 µL. Each reaction contained 1 µL of 10× Tris–Mg buffer (300 mM MgCl_2_ + 200 mM Tris-HCl), 2 µL of purified nanoswitch (∼400 pM), and 1 µL of 1 nM DNA target or nuclease-free water for control reactions, along with 6.5 µL of NFW to adjust the final volume. The mixtures were gently mixed and incubated under the appropriate experimental conditions as required for the ligation process.

### Nanoswitch reactions with ligase

The ligation reactions were prepared in a total volume of 10 µL containing 1 µL of ligase buffer, 2 µL of nanoswitch (V8 5′-phosphorylated), and 1 µL of Let-7b DNA target (variable concentration) or nuclease-free water (NFW) for control reactions. An additional 5 µL of NFW was added to adjust the reaction volume, followed by the addition of 1 µL of ligase (variable U/µL). The reaction mixtures were gently mixed and incubated under the appropriate ligation conditions prior to downstream analysis.

### Gel Electrophoresis

Reaction products were analyzed by agarose gel electrophoresis using 0.8% agarose gels prepared and run in 0.5x TBE buffer. Electrophoresis was performed at 75 V for 45 minutes, after which the gels were stained in 1x GelRed DNA stain prepared in 0.5x TBE and visualized to assess product formation and looping efficiency.

## Acknowledgements

The authors acknowledge Arun Richard Chandrasekaran for useful discussions, and Lillie Carnell for early experiments in this area. Research reported in this publication was supported by the National Institutes of Health (NIH) through National Institute of General Medical Sciences (NIGMS) under award number R35GM124720 (to K.H.). This manuscript is the result of funding in whole or in part by the National Institutes of Health (NIH). It is subject to the NIH Public Access Policy. Through acceptance of this federal funding, NIH has been given a right to make this manuscript publicly available in PubMed Central upon the Official Date of Publication, as defined by NIH.

