## Supplemental Information for "Ligation-assisted target recycling for DNA nanoswitch biosensors"

12561 USA.

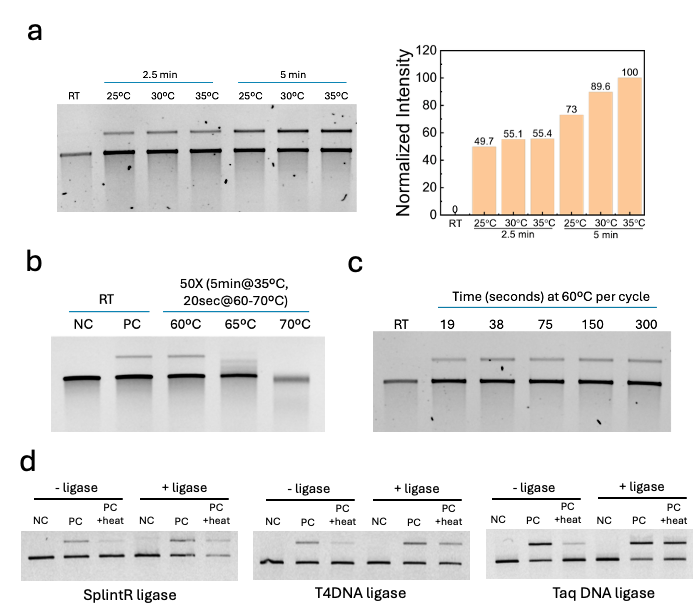

Figure S1. Optimization of a) annealing temperature and time required for looping, b) temperature required for target removal, c) time required for target removal.

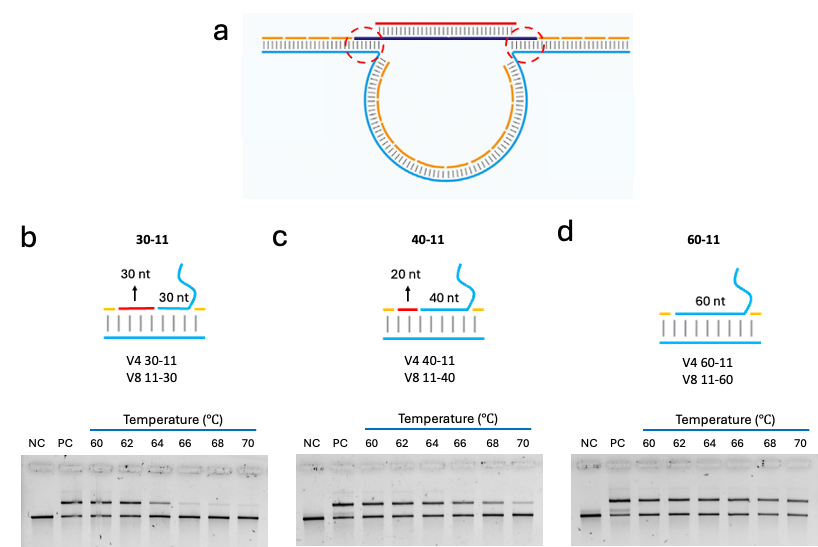

Figure S2. Comparison of different backbone oligo length on the detector oligos. a) Schematic of the region of interest. Samples were incubated for 10 minutes after ligation at different temperature with b) 30nt long, c) 40nt long and d) 60nt long backbone region.

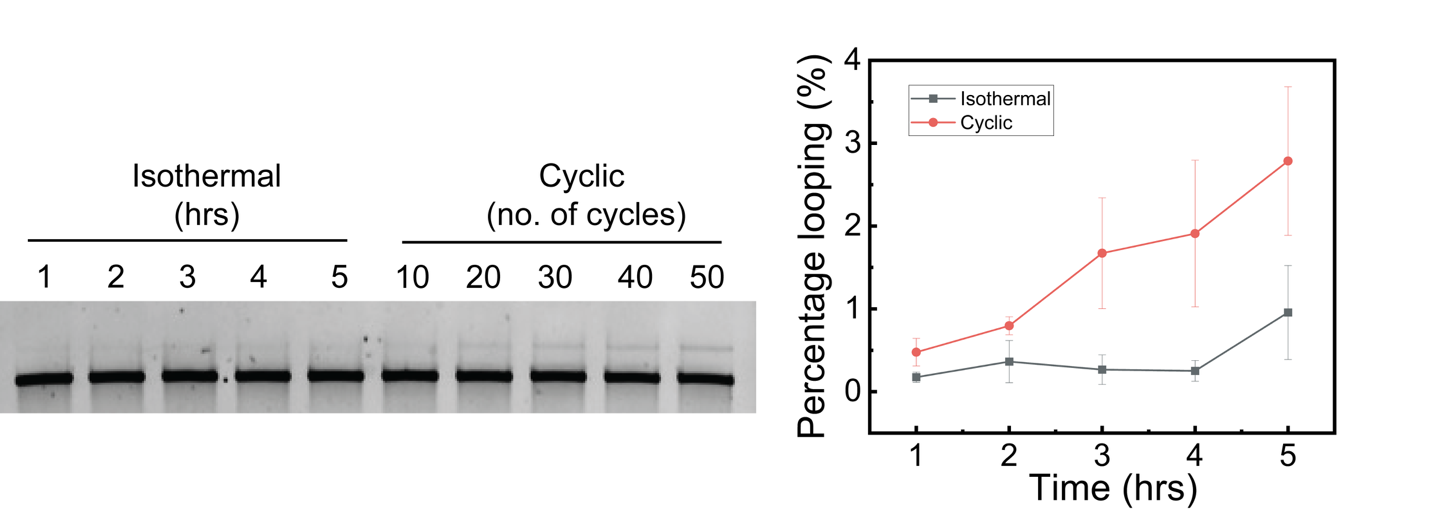

Figure S3. Comparison of isothermal vs cyclic ligation reactions. The 10, 20, 30, 40 and 50 cycles take around 1, 2, 3, 4 and 5 hours, respectively.

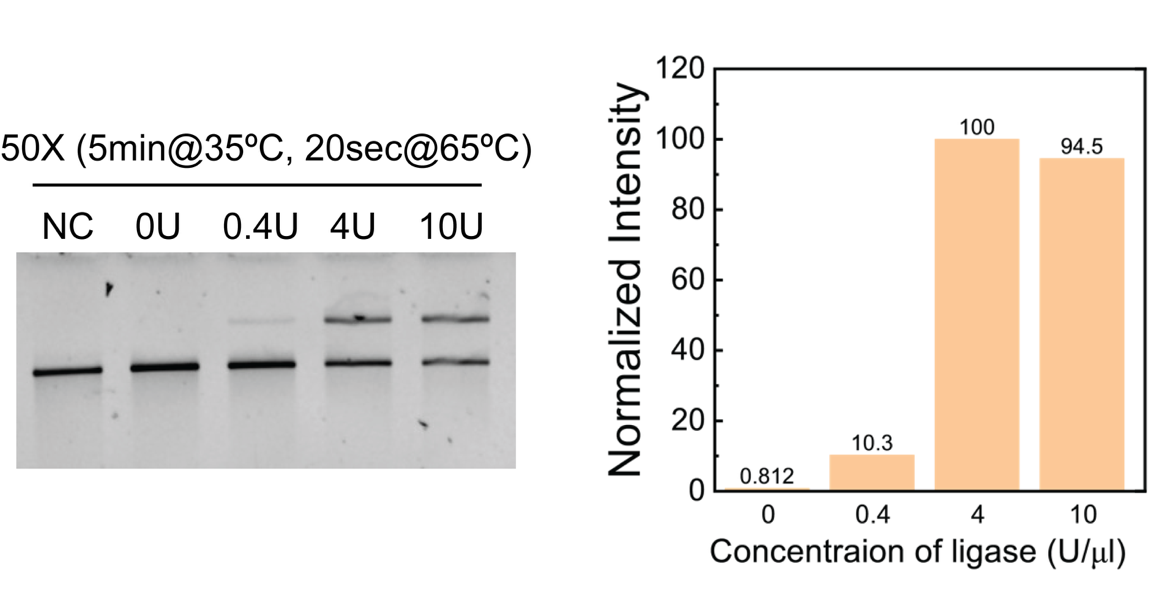

Figure S4. Effect of ligase concentration on the looping efficiency.

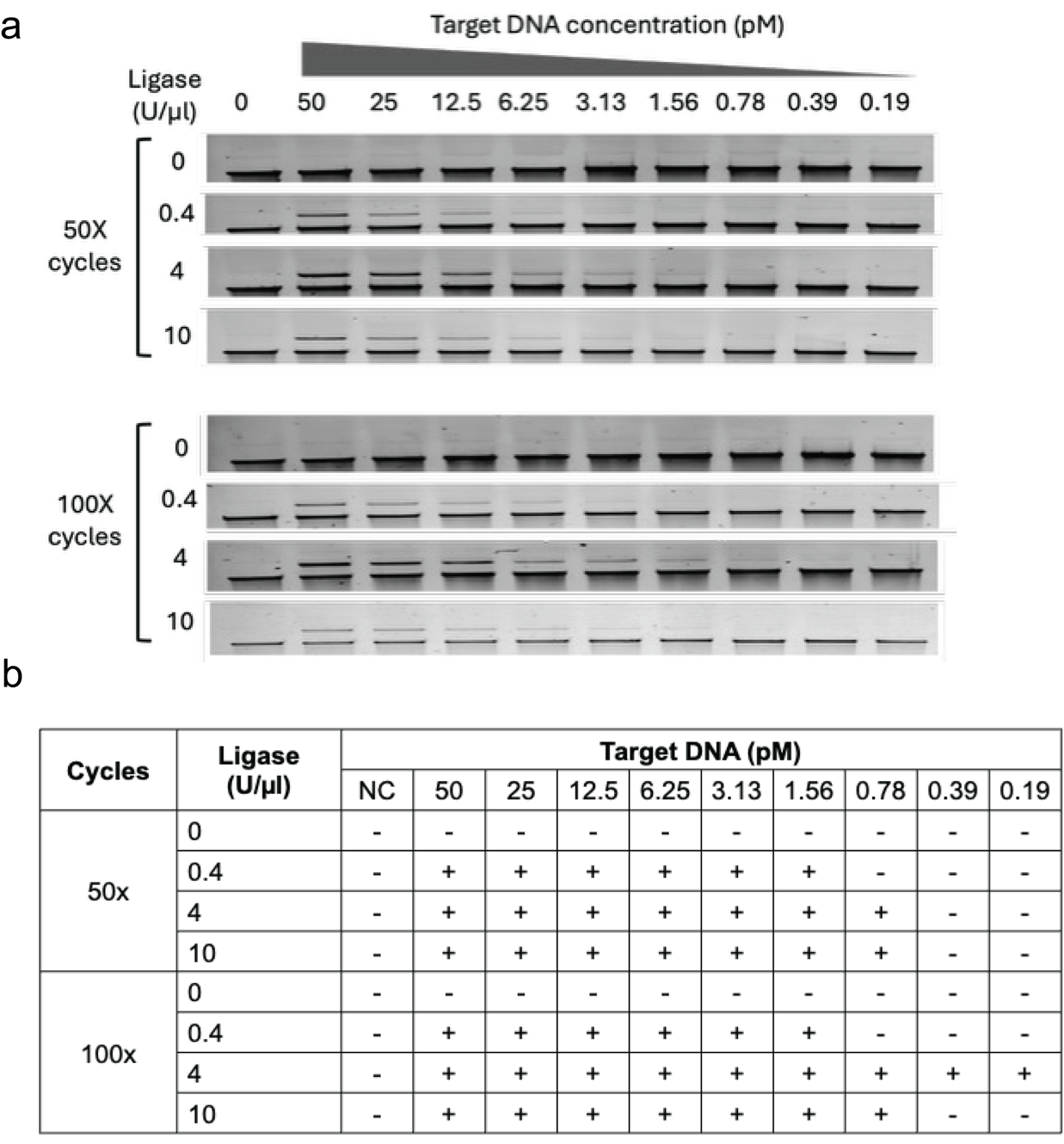

Figure S5. a) Sensitivity test for let-7b DNA with increasing concentrations of ligase for 50 and 100 cycles of reactions. b) Visual observation record from panel (a). Where - = no visible band, + = visible band.
